## Supplementary material for "Myoglobin primary structure reveals multiple convergent transitions to semi-aquatic life in the world’s smallest mammalian divers": text S1

**Supplementary Text 1. Calibrations used in divergence time estimation in BEAST2.**

1. **Boreoeutheria** (The crown group of all taxa, using *Cavia porcellus* as outgroup)

**Type**: A secondary Calibration.

**Calibration: Uniform Distribution, 83-107 million years (Ma)**

**Discussion.** The common ancestor of Euarchontoglires and Laurasiatheria. Fossil record of the crown Placentalia is not very informative and could only calibrate the K/Pg boundary (61.7 Ma ± 0.1 Ma). We calibrated it based on the divergence time estimated in (Meredith, et al. 2011). The time estimated in (Meredith, et al. 2011) was 92.0 (82.9–107.6) Ma.

2. **Laurasiatheria**

**Type**: A secondary Calibration.

**Calibration: Normal** distribution. **Mean: 85.75, sigma 4.4 Ma,** so the 95%CI is 78.5-93 Ma.

**Discussion**: Fossil record of the crown Laurasiatheria is not very informative and could only calibrate the K/Pg boundary (61.6 Ma). We calibrated it based on (Meredith, et al. 2011), which was at 84.6 (78.5–93.0) Ma.

3. **Eulipotyphla (The crown group of Erinaceidae + Soricidae + Talpidae + Solenodontidae)**

**Type**: A secondary Calibration.

**Calibration:** A **normal** distribution was used for this calibration with a mean of **78.3 Ma and a sigma of 4.6 Ma**, and the 95%CI is 70.7-85.9 Ma.

**Discussion.** Based on the results of (Meredith, et al. 2011), the 95% confidence intervals of Eulipotyphla are 70.7 to 85.8 Ma.

4. **Most common recent ancestor (MCRA) of Soricidae + Erinaceidae + Talpidae**

**Type**: A secondary Calibration.

**Calibration:** A **normal** distribution was used for this calibration with a mean of **75.76 Ma and sigma of 5.5 Ma**, and the 95%CI is 66.7-84.8 Ma.

**Discussion.** Based on the results of (Meredith, et al. 2011), the 95% confidence intervals are 66.7 to 84.8 Ma for this clade.

5. **Stem Talpini**

**Type**: A fossil-based Calibration.

**Fossil Taxon and Specimen.** *Geotrypus minor* (SMNS 47462; A1) from Ehrenstein 12, a fissure filling in a quarry in Ehrenstein, a sub-municipality of the city Blaustein near Ulm (Ziegler 2012).

**Minimum Age.** 33.9 Ma (İslamoğlu, et al. 2010). All samples correlate with the Early Oligocene standard levels of Soumailles (MP 21) and Villebramar (MP 22) (Ziegler 2012).

**Calibration:** An **exponential distribution** was used for this calibration with offset of **33.9 Ma and mean of 11.3 Ma** (median=41.73 Ma, 95%CI=34.48 to 67.75 Ma).
**Discussion.** *Geotrypus minor* is the oldest record of the genus and the smallest species of *Geotrypus* (Ziegler 2012). The genus was initially included in Scaptonychini, but later referred to Talpini (Ziegler 1990; van den Hoek Ostende 2001) because of their robust humeri. However, the overall shape of the humerus of *G. minor* is more similar to a shrew mole than to a true mole. It is more appropriate to place it on the stem rather than the crown of living Talpini. There is no reliable upper boundary for calibration, so we follow (Heath 2012), in which suggested "in the absence of prior knowledge of the true node to fossil age differences, it is advisable to assign a vague prior distribution to calibrated node ages", i.e. to use exponential distribution with a mean value of: fossil age * 0.333.
*#note*: this calibration is the same as calibration "10 Stem Talpini" in the Supplementary Text S4 in (He, et al. 2017).

**6. MCRA of Soricidae**
**Type**: A secondary Calibration.
**Calibration**: A normal distribution. We set the mean **to 36 Ma, with standard deviation to 0.135 Ma, so that the median of prior is 35.7 Ma a**nd 95% CI is 28.6-44.5 Ma.
**Discussion**: We applied a second calibration following (Springer, et al. 2018), which focused on the divergence times of eulipotyphlans and estimated that the divergence between Crocidurinae and Soricinae occurred 36 million years ago (Ma) [95% confidence interval (CI) = 28.6-44.0 Ma]. This time is much older than a fossil calibration used in previous studies (Dubey, et al. 2007; He, et al. 2010), which was based on the assumption that Crocidosoricinae was the ancestor of Crocidurinae, Myosoricinae, and Soricinae. Crocidosoricinae is now recognized as a tribe of Myosoricinae (Crocidosoricini), thereby invalidating that assumption (Furio, et al. 2007). Fossils of both crocidurines and soricines are known from the Oligocene (< 34 Ma), and the oldest soricid fossil, *Soricolestes soricavus* is from Middle Eocene strata, Khaychin Formation (Lopatin 2002). Thus, the estimated divergence time in (Springer, et al. 2018) is congruent with fossil records.

**7. MCRA of Anourosoricini and Nectogalini**
**Type**: A fossil-based Calibration.
**Fossil Taxon and Specimen.** *Darocasorex vandermeuleni* (Naturalis, Leiden: 558887: left M1;558857: right m2; 558855: left m3) from Calatayud-Daroca Basin, east Central Spain. Level at transition local biozone G to H (MN 7/8 to 9), ~11.5 Ma (van Dam 2010).
**Minimum Age.** 12 Ma (van Dam 2010).
**Calibration.** An exponential distribution for the prior of this calibration with offset of 12 Ma and a mean of 4 Ma (median=14.8 Ma, 95%CI=12.2-24.0 Ma).
**Discussion.** Anourosoricini is a fossil rich group. In the Late Miocene at about 12-10 Ma,

*Darocasorex* and *Crusafontina* (Anourosoricini) already existed (van Dam 2010). The oldest fossil of Nectogalini represented by *Asoriculus* (Nectogalini) from Europe at about MN10 (=8.7-9.7 Ma)(Fejfar and Sabol 2005; Rzebik-Kowalska and Lungu 2009). Thus, it is convincing that Anourosoricini diverged from Nectogalini and Notiosoricini more than 12 million years ago. We set the lower boundary to 12 million years ago. There is no reliable upper boundary for calibration, so we used exponential distribution with a mean value of: fossil age * 0.333.

van den Hoek Ostende LW. 2001. Insectivore faunas from the Lower Miocene of Anatolia-Part 8:

Stratigraphy, palaeoecology, palaeobiogeography. Scripta Geologica 122:101-122.
Ziegler R. 2012. Moles (Talpidae, Mammalia) from Early Oligocene karstic fissure fillings in South Germany. Geobios 45:501-513.
Ziegler R. 1990. Talpidae (Insectivora, Mammalia) aus dem Oberoligozan und Untermiozan Suddeutschlands. Stuttgarter Beitraege zur Naturkunde Serie B (Geologie und Palaeontologie) 167:1-81.
