## Supplementary material for "Myoglobin primary structure reveals multiple convergent transitions to semi-aquatic life in the world’s smallest mammalian divers": fig. S

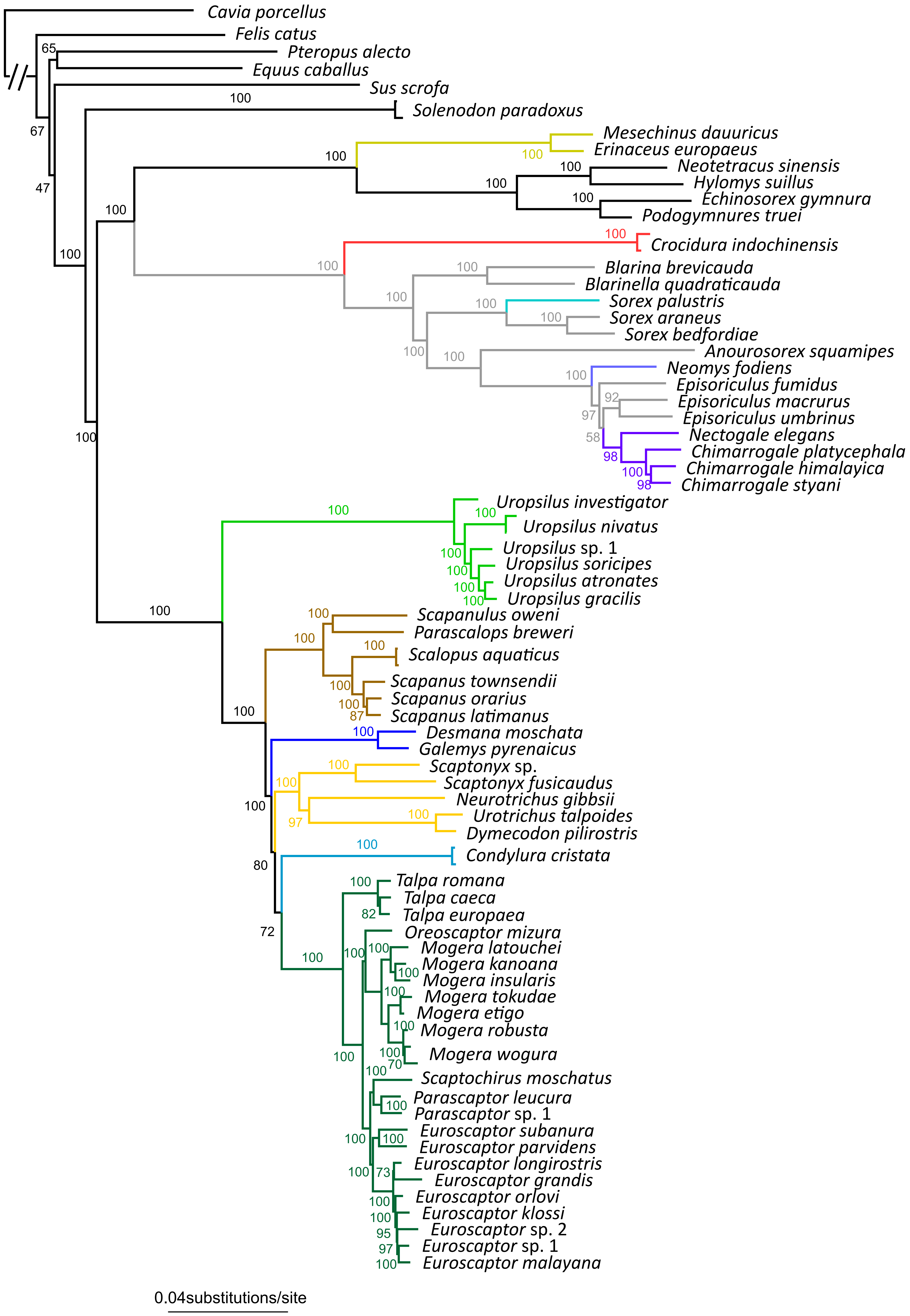
Cavia porcellus
Felis catus
Pteropus alecto
Equus caballus
Sus scrofa
Solenodon paradoxus
Mesechinus dauuricus
Erinaceus europaeus
Neotetracus sinensis
Hylomys suillus
Echinosorex gymnura
Podogymnures truei
Crocidura indochinensis
Blarina brevicauda
Blarinella quadraticauda
Sorex palustris
Sorex araneus
Sorex bedfordiae
Anourosorex squamipes
Neomys fodiens
Episoriculus fumidus
Episoriculus macrurus
Episoriculus umbrinus
Nectogale elegans
Chimarrogale platycephala
Chimarrogale himalayica
Chimarrogale styani
Uropsilus investigator
Uropsilus nivatus
Uropsilus sp. 1
Uropsilus soricipes
Uropsilus atronates
Uropsilus gracilis
Scapanulus oweni
Parascalops breweri
Scalopus aquaticus
Scapanus townsendii
Scapanus orarius
Scapanus latimanus
Desmana moschata
Galemys pyrenaicus
Scaptonyx sp.
Scaptonyx fusicaudus
Neurotrichus gibbsii
Urotrichus talpoides
Dymecodon pilirostris
Condylura cristata
Talpa romana
Talpa caeca
Talpa europaea
Oreoscaptor mizura
Mogera latouchei
Mogera kanoana
Mogera insularis
Mogera tokudae
Mogera etigo
Mogera robusta
Mogera wogura
Scaptochirus moschatus
Parascaptor leucura
Parascaptor sp. 1
Euroscaptor subanura
Euroscaptor parvidens
Euroscaptor longirostris
Euroscaptor grandis
Euroscaptor orlovi
Euroscaptor klossi
Euroscaptor sp. 2
Euroscaptor sp. 1
Euroscaptor malayana
0.04substitutions/site

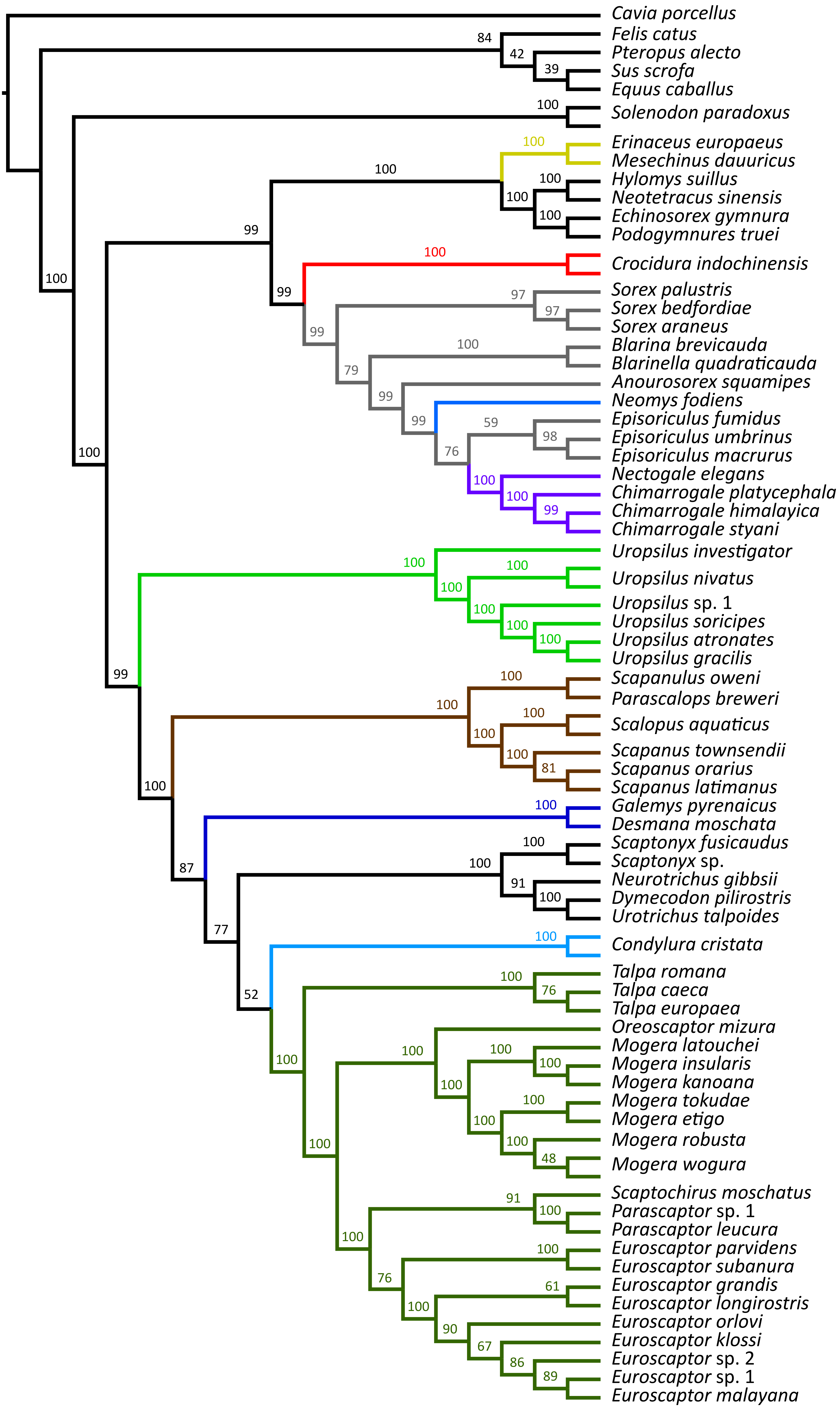
Cavia porcellus
84
Felis catus
42
Pteropus alecto
39
Sus scrofa
Equus caballus
100
Solenodon paradoxus
100
Erinaceus europaeus
Mesechinus dauuricus
100
100
Hylomys suillus
100
Neotetracus sinensis
100
Echinosorex gymnura
Podogymnures truei
99
100
Crocidura indochinensis
100
99
97
Sorex palustris
97
Sorex bedfordiae
Sorex araneus
99
100
Blarina brevicauda
Blarinella quadraticauda
79
Anourosorex squamipes
99
Neomys fodiens
59
Episoriculus fumidus
99
98
Episoriculus umbrinus
76
Episoriculus macrurus
Nectogale elegans
100
100
Chimarrogale platycephala
99
Chimarrogale himalayica
Chimarrogale styani
100
Uropsilus investigator
100
100
Uropsilus nivatus
100
Uropsilus sp. 1
100
Uropsilus soricipes
100
Uropsilus atronates
Uropsilus gracilis
99
100
Scapanulus oweni
100
Parascalops breweri
100
Scalopus aquaticus
100
Scapanus townsendii
100
81
Scapanus orarius
Scapanus latimanus
100
100
Galemys pyrenaicus
Desmana moschata
100
Scaptonyx fusicaudus
100
87
Scaptonyx sp.
91
Neurotrichus gibbsii
100
Dymecodon pilirostris
77
Urotrichus talpoides
100
Condylura cristata
100
Talpa romana
76
Talpa caeca
52
Talpa europaea
Oreoscaptor mizura
100
Mogera latouchei
100
100
Mogera insularis
100
100
Mogera kanoana
100
Mogera tokudae
100
Mogera etigo
100
100
Mogera robusta
48
Mogera wogura
Scaptochirus moschatus
91
Parascaptor sp. 1
100
Parascaptor leucura
100
100
Euroscaptor parvidens
Euroscaptor subanura
76
61
Euroscaptor grandis
100
Euroscaptor longirostris
Euroscaptor orlovi
90
Euroscaptor klossi
67
Euroscaptor sp. 2
86
Euroscaptor sp. 1
89
Euroscaptor malayana

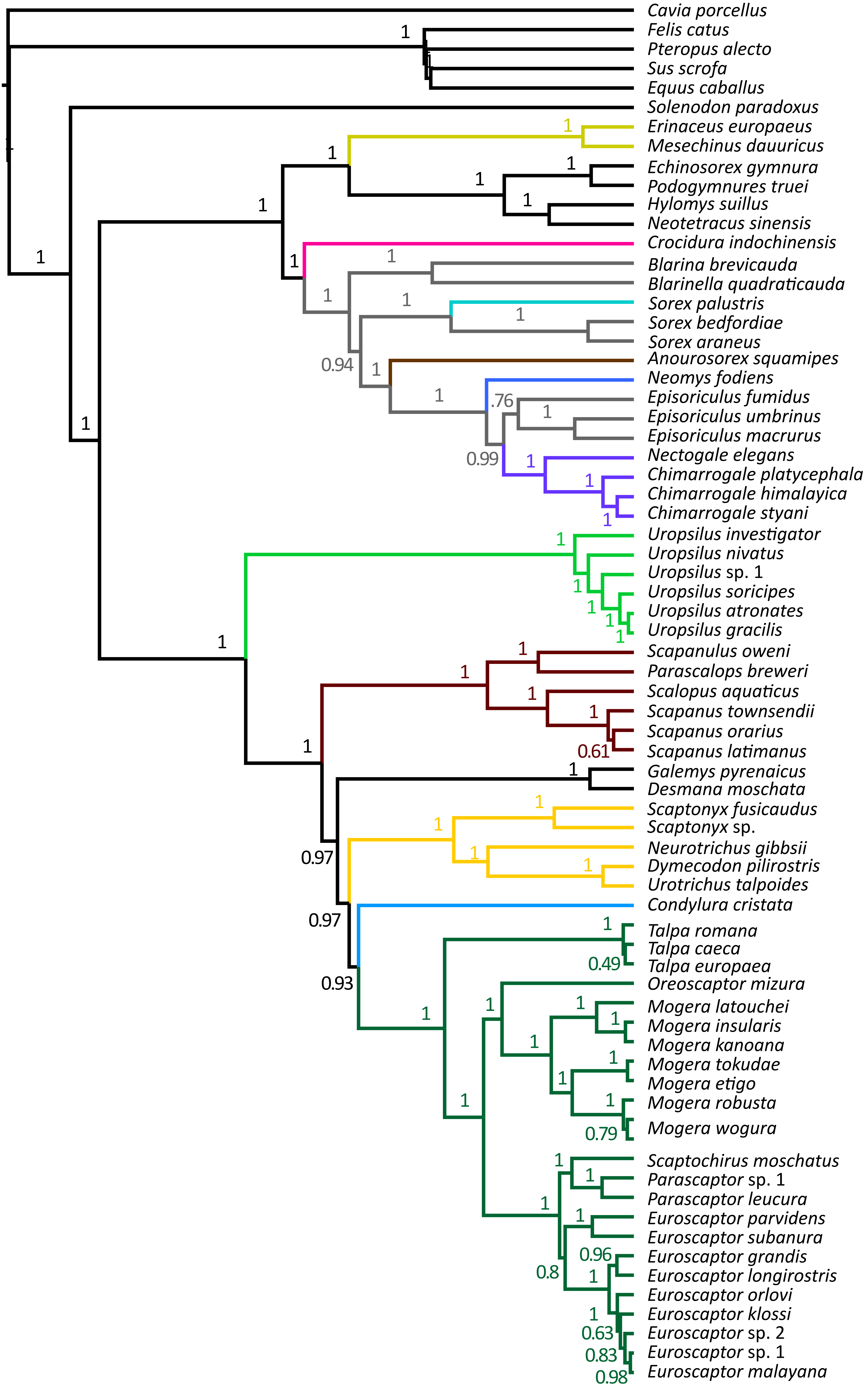
Cavia porcellus
Felis catus
Pteropus alecto
Sus scrofa
Equus caballus
Solenodon paradoxus
Erinaceus europaeus
Mesechinus dauuricus
Echinosorex gymnura
Podogymnures truei
Hylomys suillus
Neotetracus sinensis
Crocidura indochinensis
Blarina brevicauda
Blarinella quadraticauda
Sorex palustris
Sorex bedfordiae
Sorex araneus
Anourosorex squamipes
Neomys fodiens
Episoriculus fumidus
Episoriculus umbrinus
Episoriculus macrurus
Nectogale elegans
Chimarrogale platycephala
Chimarrogale himalayica
Chimarrogale styani
Uropsilus investigator
Uropsilus nivatus
Uropsilus sp. 1
Uropsilus soricipes
Uropsilus atronates
Uropsilus gracilis
Scapanulus oweni
Parascalops breweri
Scalopus aquaticus
Scapanus townsendii
Scapanus orarius
Scapanus latimanus
Galemys pyrenaicus
Desmana moschata
Scaptonyx fusicaudus
Scaptonyx sp.
Neurotrichus gibbsii
Dymecodon pilirostris
Urotrichus talpoides
Condylura cristata
Talpa romana
Talpa caeca
Talpa europaea
Oreoscaptor mizura
Mogera latouchei
Mogera insularis
Mogera kanoana
Mogera tokudae
Mogera etigo
Mogera robusta
Mogera wogura
Scaptochirus moschatus
Parascaptor sp. 1
Parascaptor leucura
Euroscaptor parvidens
Euroscaptor subanura
Euroscaptor grandis
Euroscaptor longirostris
Euroscaptor orlovi
Euroscaptor klossi
Euroscaptor sp. 2
Euroscaptor sp. 1
Euroscaptor malayana

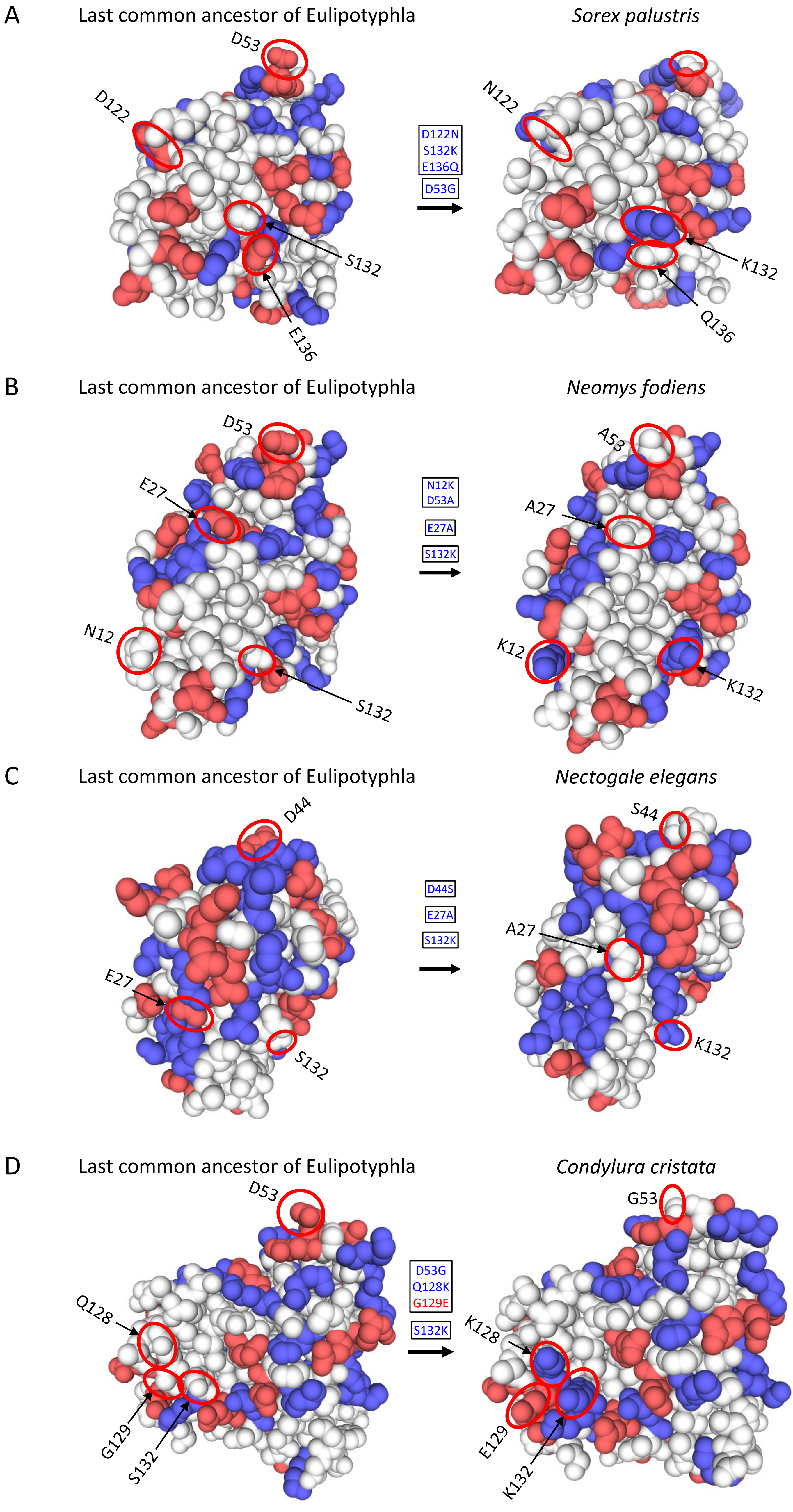
A
Last common ancestor of Eulipotyphla
Sorex palustris
D53
D122
S132
E136
D122N
S132K
E136Q
D53G
N122
K132
Q136
B
Last common ancestor of Eulipotyphla
Neomys fodiens
D53
E27
N12
S132
N12K
D53A
E27A
S132K
A53
A27
K12
K132
C
Last common ancestor of Eulipotyphla
Nectogale elegans
D44
E27
S132
D44S
E27A
S132K
S44
A27
K132
D
Last common ancestor of Eulipotyphla
Condylura cristata
D53
Q128
G129
S132
D53G
Q128K
G129E
S132K
G53
K128
E129
K132

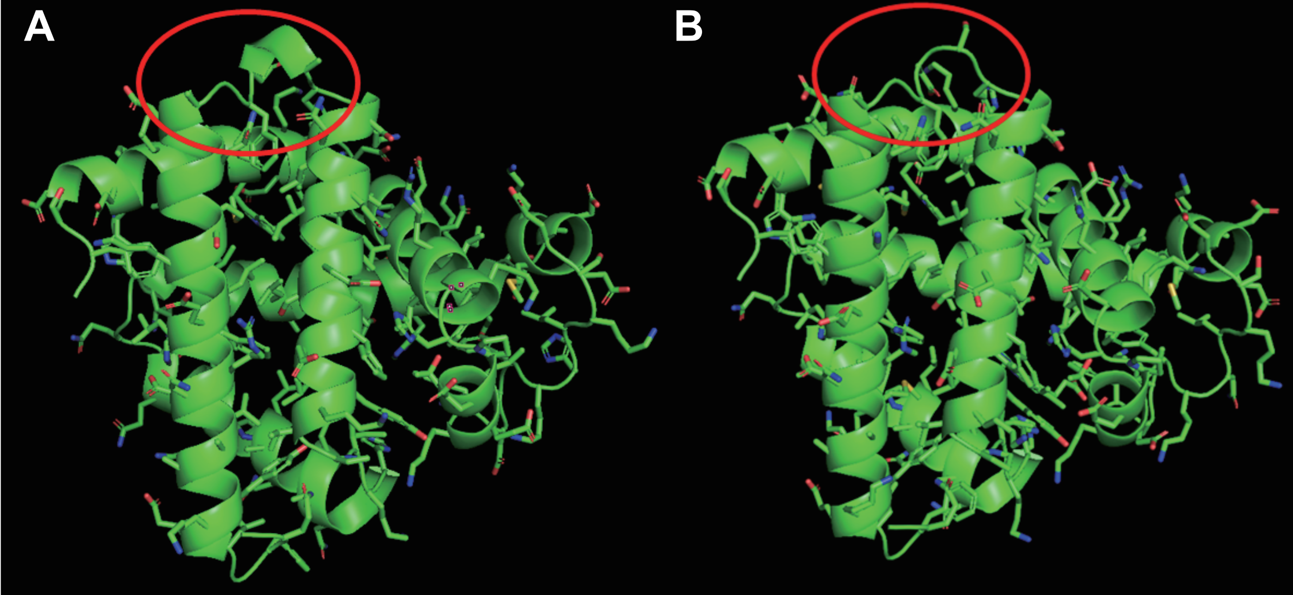
A
B

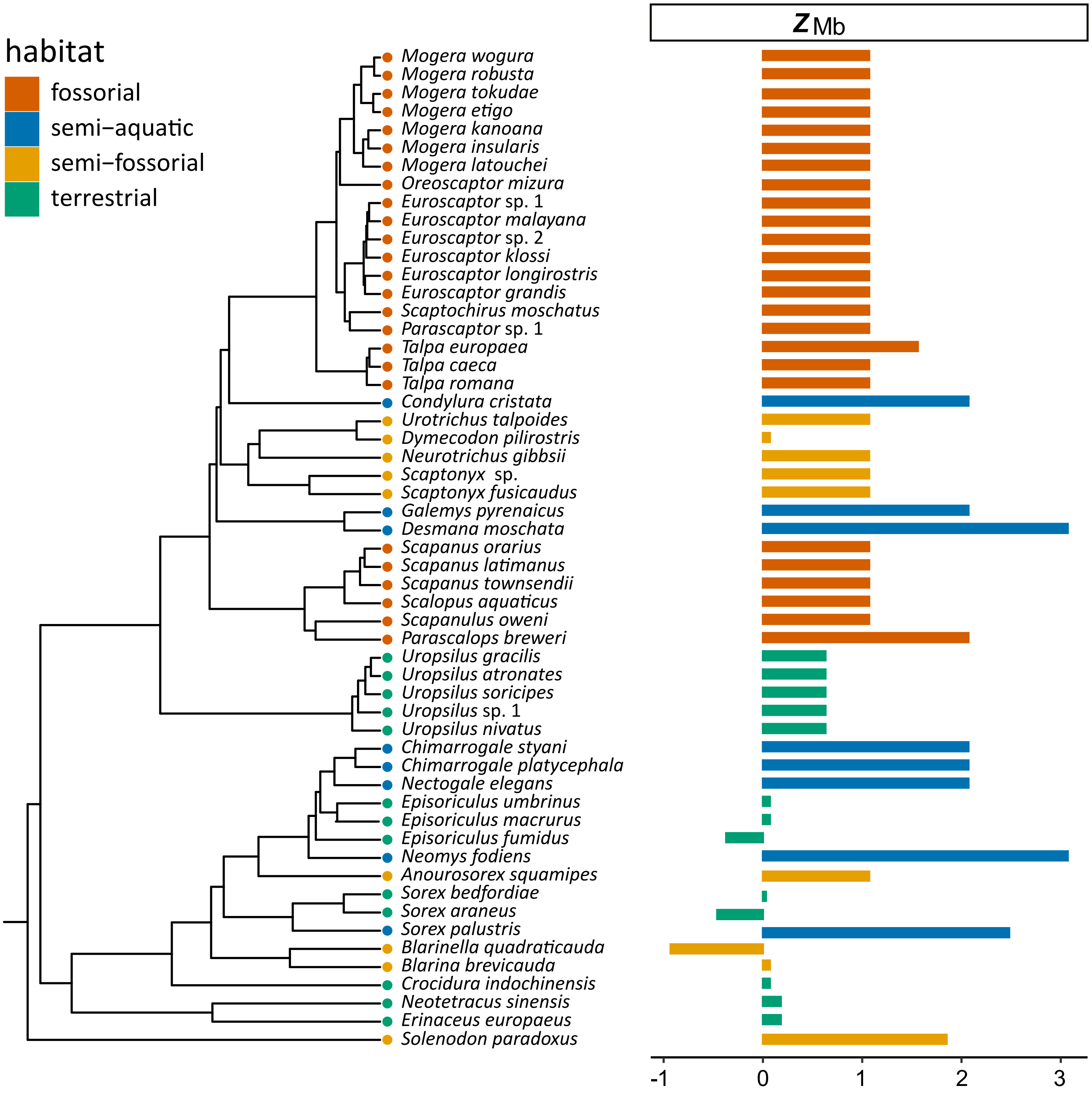
habitat
fossorial
semi−aquatic
semi−fossorial
terrestrial
*Mogera wogura*
*Mogera robusta*
*Mogera tokudae*
*Mogera etigo*
*Mogera kanoana*
*Mogera insularis*
*Mogera latouchei*
*Oreoscaptor mizura*
*Euroscaptor* sp. 1
*Euroscaptor malayana*
*Euroscaptor* sp. 2
*Euroscaptor klossi*
*Euroscaptor longirostris*
*Euroscaptor grandis*
*Scaptochirus moschatus*
*Parascaptor* sp. 1
*Talpa europaea*
*Talpa caeca*
*Talpa romana*
*Condylura cristata*
*Urotrichus talpoides*
*Dymecodon pilirostris*
*Neurotrichus gibbsii*
*Scaptonyx* sp.
*Scaptonyx fusicaudus*
*Galemys pyrenaicus*
*Desmana moschata*
*Scapanus orarius*
*Scapanus latimanus*
*Scapanus townsendii*
*Scalopus aquaticus*
*Scapanulus oweni*
*Parascalops breweri*
*Uropsilus gracilis*
*Uropsilus atronates*
*Uropsilus soricipes*
*Uropsilus* sp. 1
*Uropsilus nivatus*
*Chimarrogale styani*
*Chimarrogale platycephala*
*Nectogale elegans*
*Episoriculus umbrinus*
*Episoriculus macrurus*
*Episoriculus fumidus*
*Neomys fodiens*
*Anourosorex squamipes*
*Sorex bedfordiae*
*Sorex araneus*
*Sorex palustris*
*Blarinella quadraticauda*
*Blarina brevicauda*
*Crocidura indochinensis*
*Neotetracus sinensis*
*Erinaceus europaeus*
*Solenodon paradoxus*
$z_{Mb}$
-1
0
1
2
3

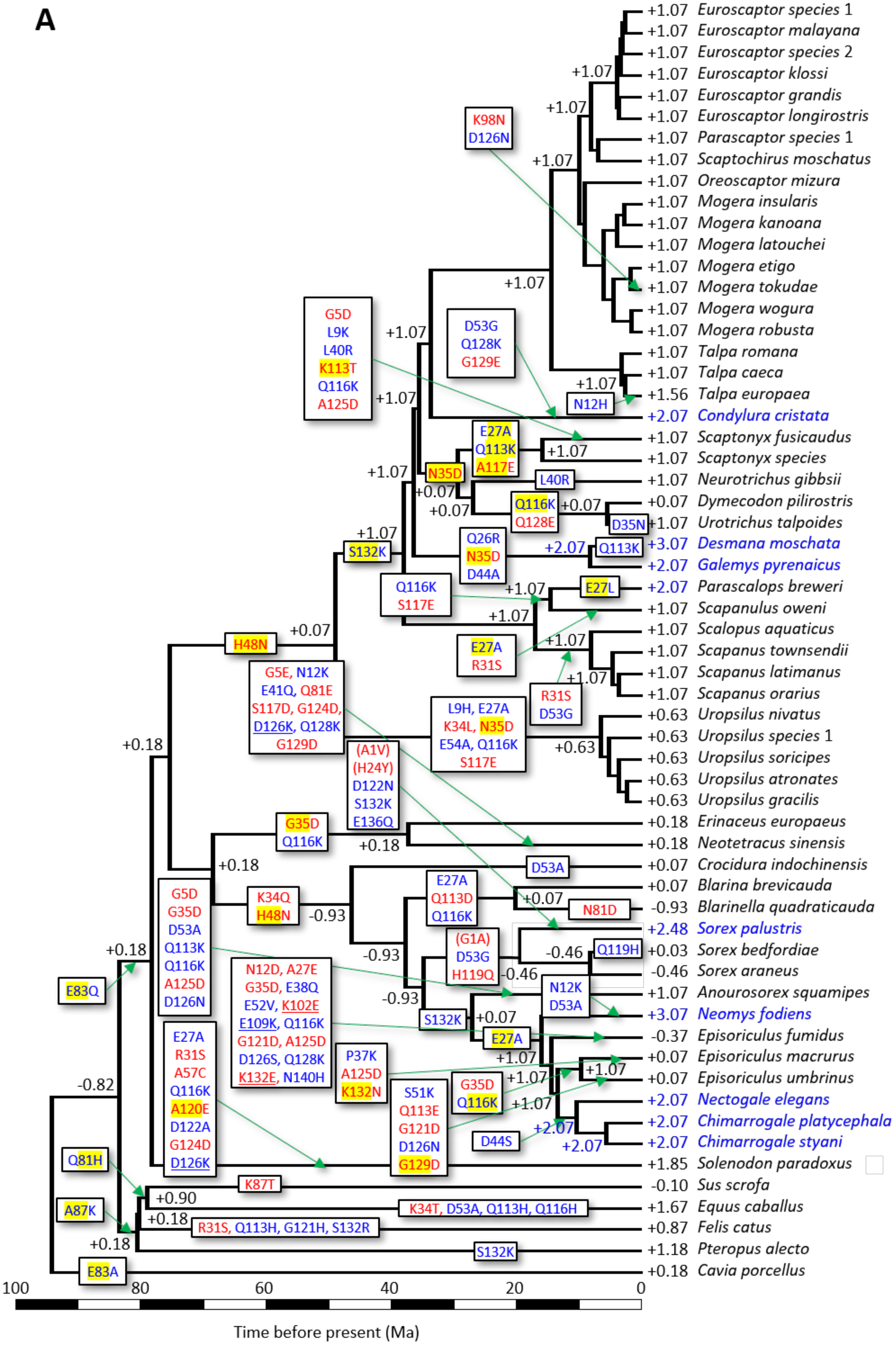

A
+1.07 Euroscaptor species 1
+1.07 Euroscaptor malayana
+1.07 Euroscaptor species 2
+1.07 Euroscaptor klossi
+1.07 Euroscaptor grandis
+1.07 Euroscaptor longirostris
+1.07 Parascaptor species 1
+1.07 Scaptochirus moschatus
+1.07 Oreoscaptor mizura
+1.07 Mogera insularis
+1.07 Mogera kanoana
+1.07 Mogera latouchei
+1.07 Mogera etigo
+1.07 Mogera tokudae
+1.07 Mogera wogura
+1.07 Mogera robusta
+1.07 Talpa romana
+1.07 Talpa caeca
+1.56 Talpa europaea
+2.07 Condylura cristata
+1.07 Scaptonyx fusicaudus
+1.07 Scaptonyx species
+1.07 Neurotrichus gibbsii
+0.07 Dymecodon pilirostris
+1.07 Urotrichus talpoides
+3.07 Desmana moschata
+2.07 Galemys pyrenaicus
+2.07 Parascalops breweri
+1.07 Scapanulus oweni
+1.07 Scalopus aquaticus
+1.07 Scapanus townsendii
+1.07 Scapanus latimanus
+1.07 Scapanus orarius
+0.63 Uropsilus nivatus
+0.63 Uropsilus species 1
+0.63 Uropsilus soricipes
+0.63 Uropsilus atronates
+0.63 Uropsilus gracilis
+0.18 Erinaceus europaeus
+0.18 Neotetracus sinensis
+0.07 Crocidura indochinensis
+0.07 Blarina brevicauda
-0.93 Blarinella quadraticauda
+2.48 Sorex palustris
+0.03 Sorex bedfordiae
-0.46 Sorex araneus
+1.07 Anourosorex squamipes
+3.07 Neomys fodiens
-0.37 Episoriculus fumidus
+0.07 Episoriculus macrurus
+0.07 Episoriculus umbrinus
+2.07 Nectogale elegans
+2.07 Chimarrogale platycephala
+2.07 Chimarrogale styani
+1.85 Solenodon paradoxus
-0.10 Sus scrofa
+1.67 Equus caballus
+0.87 Felis catus
+1.18 Pteropus alecto
+0.18 Cavia porcellus
K98N
D126N
G5D
L9K
L40R
K113T
Q116K
A125D
D53G
Q128K
G129E
N12H
E27A
Q113K
A117E
N35D
L40R
Q116K
Q128E
D35N
Q26R
N35D
D44A
S132K
Q116K
S117E
E27L
H48N
E27A
R31S
R31S
D53G
G5E, N12K
E41Q, Q81E
S117D, G124D,
D126K, Q128K
G129D
L9H, E27A
K34L, N35D
E54A, Q116K
S117E
(A1V)
(H24Y)
D122N
S132K
E136Q
G35D
Q116K
D53A
K34Q
H48N
E27A
Q113D
Q116K
N81D
G5D
G35D
D53A
Q113K
Q116K
A125D
D126N
(G1A)
D53G
H119Q
Q119H
N12K
D53A
E83Q
N12D, A27E
G35D, E38Q
E52V, K102E
E109K, Q116K
G121D, A125D
D126S, Q128K
K132E, N140H
S132K
E27A
E27A
R31S
A57C
Q116K
A120E
D122A
G124D
D126K
P37K
A125D
K132N
S51K
Q113E
G121D
D126N
G129D
G35D
Q116K
D44S
Q81H
K87T
A87K
K34T, D53A, Q113H, Q116H
R31S, Q113H, G121H, S132R
S132K
E83A
+1.07
+1.07
+1.07
+1.07
+1.07
+1.07
+1.07
+1.07
+0.07
+0.07
+0.07
+2.07
+1.07
+1.07
+1.07
+1.07
+1.07
+0.07
+0.63
+0.18
+0.18
+0.18
-0.93
-0.93
-0.93
+0.07
-0.46
-0.46
+0.07
+1.07
+1.07
+1.07
+1.07
+2.07
+2.07
-0.82
+0.90
+0.18
+0.18
100
80
60
40
20
0
Time before present (Ma)

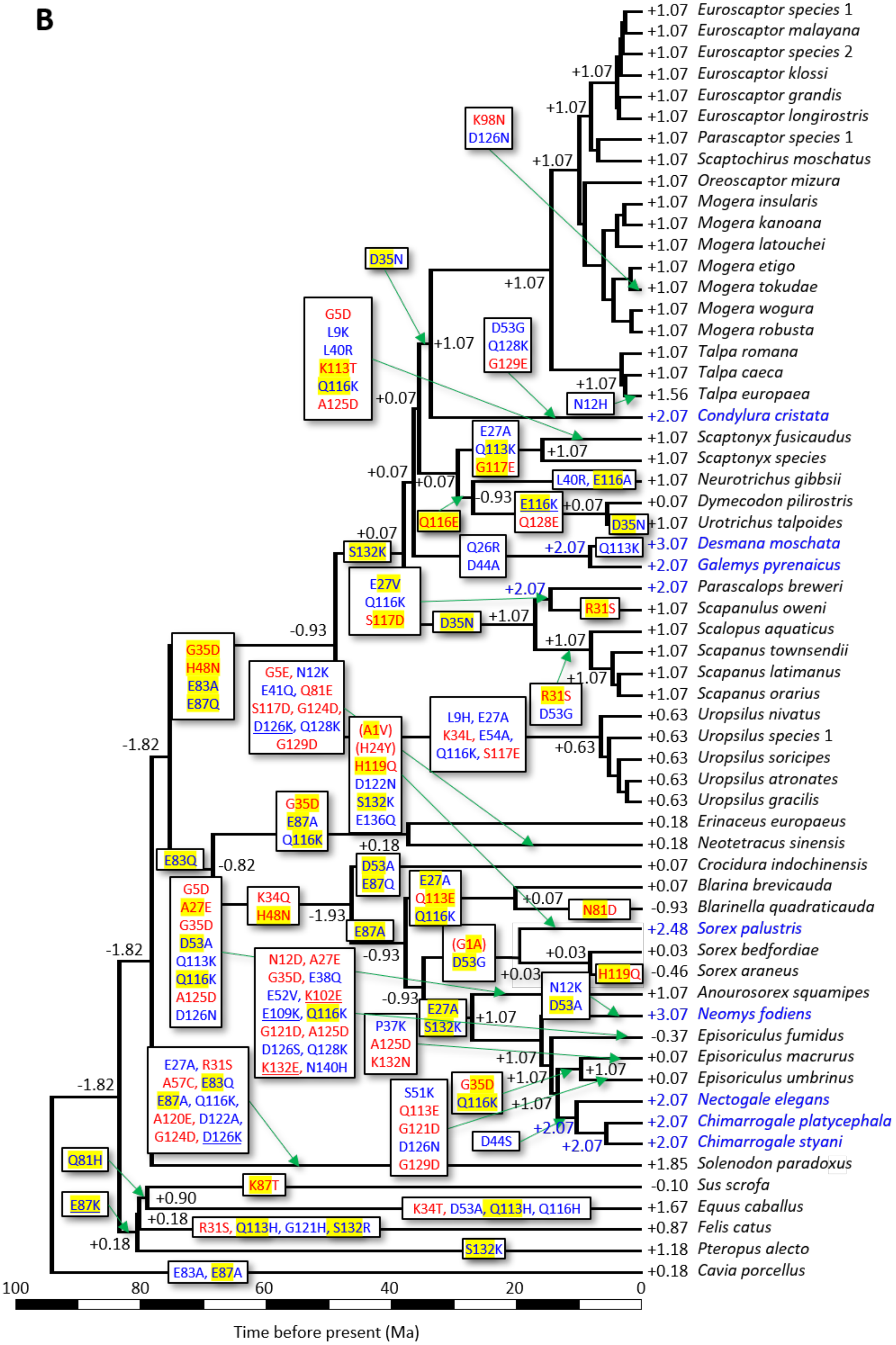
B
+1.07 Euroscaptor species 1
+1.07 Euroscaptor malayana
+1.07 Euroscaptor species 2
+1.07 Euroscaptor klossi
+1.07 Euroscaptor grandis
+1.07 Euroscaptor longirostris
+1.07 Parascaptor species 1
+1.07 Scaptochirus moschatus
+1.07 Oreoscaptor mizura
+1.07 Mogera insularis
+1.07 Mogera kanoana
+1.07 Mogera latouchei
+1.07 Mogera etigo
+1.07 Mogera tokudae
+1.07 Mogera wogura
+1.07 Mogera robusta
+1.07 Talpa romana
+1.07 Talpa caeca
+1.56 Talpa europaea
+2.07 Condylura cristata
+1.07 Scaptonyx fusicaudus
+1.07 Scaptonyx species
+1.07 Neurotrichus gibbsii
+0.07 Dymecodon pilirostris
+1.07 Urotrichus talpoides
+3.07 Desmana moschata
+2.07 Galemys pyrenaicus
+2.07 Parascalops breweri
+1.07 Scapanulus oweni
+1.07 Scalopus aquaticus
+1.07 Scapanus townsendii
+1.07 Scapanus latimanus
+1.07 Scapanus orarius
+0.63 Uropsilus nivatus
+0.63 Uropsilus species 1
+0.63 Uropsilus soricipes
+0.63 Uropsilus atronates
+0.63 Uropsilus gracilis
+0.18 Erinaceus europaeus
+0.18 Neotetracus sinensis
+0.07 Crocidura indochinensis
+0.07 Blarina brevicauda
-0.93 Blarinella quadraticauda
+2.48 Sorex palustris
+0.03 Sorex bedfordiae
-0.46 Sorex araneus
+1.07 Anourosorex squamipes
+3.07 Neomys fodiens
-0.37 Episoriculus fumidus
+0.07 Episoriculus macrurus
+0.07 Episoriculus umbrinus
+2.07 Nectogale elegans
+2.07 Chimarrogale platycephala
+2.07 Chimarrogale styani
+1.85 Solenodon paradoxus
-0.10 Sus scrofa
+1.67 Equus caballus
+0.87 Felis catus
+1.18 Pteropus alecto
+0.18 Cavia porcellus
K98N D126N
D35N
G5D L9K L40R K113T Q116K A125D
D53G Q128K G129E
N12H
E27A Q113K G117E
L40R, E116A
Q116E
E116K Q128E
D35N
S132K
Q26R D44A
Q113K
E27V Q116K S117D
D35N
R31S
R31S D53G
G35D H48N E83A E87Q
G5E, N12K E41Q, Q81E S117D, G124D, D126K, Q128K G129D
(A1V) (H24Y) H119Q D122N S132K E136Q
L9H, E27A K34L, E54A, Q116K, S117E
G35D E87A Q116K
E83Q
D53A E87Q
G5D A27E G35D D53A Q113K Q116K A125D D126N
K34Q H48N
E87A
E27A Q113E Q116K
N81D
(G1A) D53G
H119Q
N12D, A27E G35D, E38Q E52V, K102E E109K, Q116K G121D, A125D D126S, Q128K K132E, N140H
N12K D53A
E27A S132K
P37K A125D K132N
G35D Q116K
S51K Q113E G121D D126N G129D
D44S
E27A, R31S A57C, E83Q E87A, Q116K, A120E, D122A, G124D, D126K
Q81H
E87K
K87T
K34T, D53A, Q113H, Q116H
R31S, Q113H, G121H, S132R
S132K
E83A, E87A
+1.07
+0.07
-0.93
+0.93
+0.03
+0.18
-0.82
-1.93
-1.82
+0.90
+2.07
100
80
60
40
20
0
Time before present (Ma)

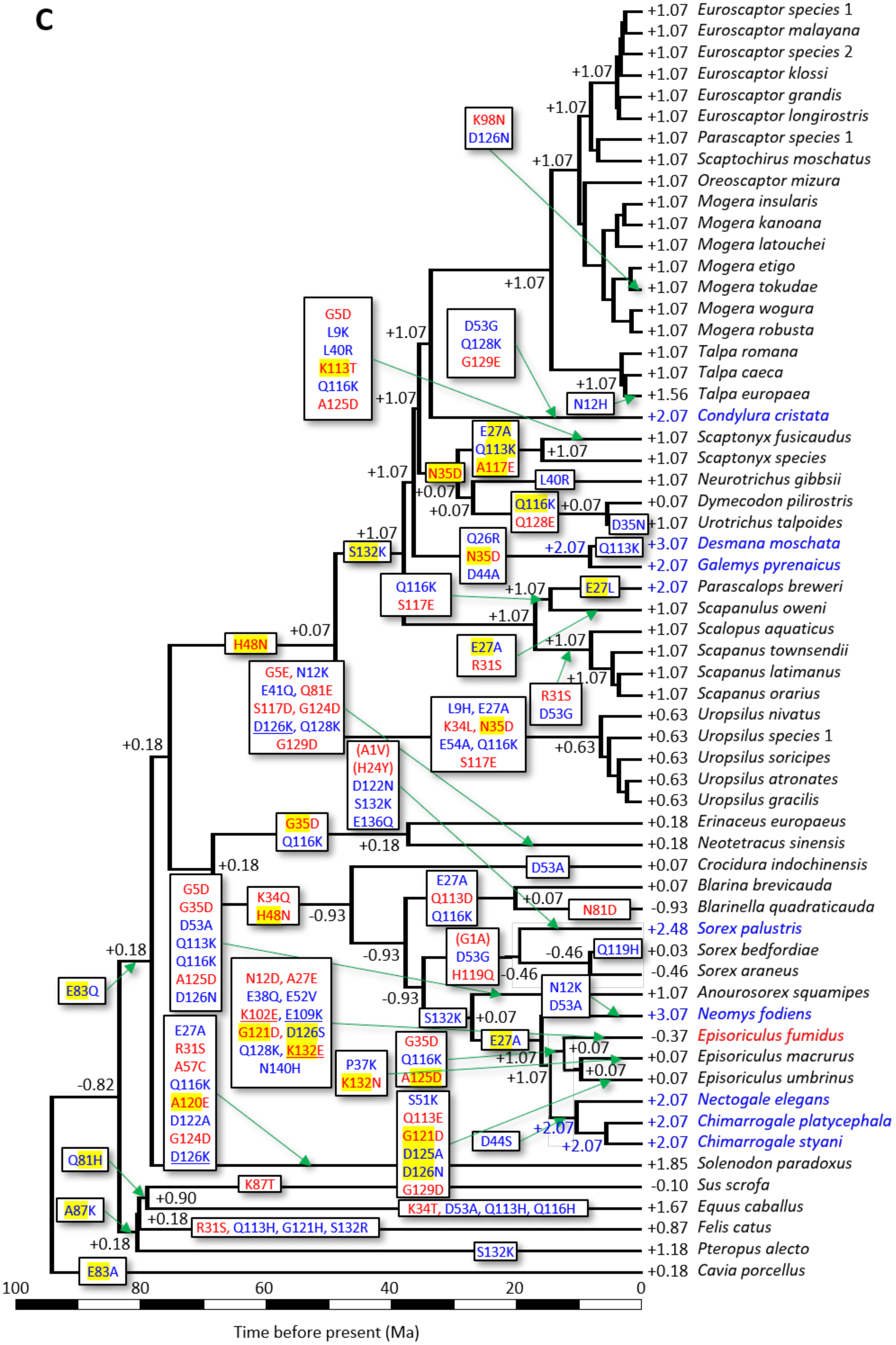

C
+1.07 Euroscaptor species 1
+1.07 Euroscaptor malayana
+1.07 Euroscaptor species 2
+1.07 Euroscaptor klossi
+1.07 Euroscaptor grandis
+1.07 Euroscaptor longirostris
+1.07 Parascaptor species 1
+1.07 Scaptochirus moschatus
+1.07 Oreoscaptor mizura
+1.07 Mogera insularis
+1.07 Mogera kanoana
+1.07 Mogera latouchei
+1.07 Mogera etigo
+1.07 Mogera tokudae
+1.07 Mogera wogura
+1.07 Mogera robusta
+1.07 Talpa romana
+1.07 Talpa caeca
+1.56 Talpa europaea
+2.07 Condylura cristata
+1.07 Scaptonyx fusicaudus
+1.07 Scaptonyx species
+1.07 Neurotrichus gibbsii
+0.07 Dymecodon pilirostris
+1.07 Urotrichus talpoides
+3.07 Desmana moschata
+2.07 Galemys pyrenaicus
+2.07 Parascalops breweri
+1.07 Scapanulus oweni
+1.07 Scalopus aquaticus
+1.07 Scapanus townsendii
+1.07 Scapanus latimanus
+1.07 Scapanus orarius
+0.63 Uropsilus nivatus
+0.63 Uropsilus species 1
+0.63 Uropsilus soricipes
+0.63 Uropsilus atronates
+0.63 Uropsilus gracilis
+0.18 Erinaceus europaeus
+0.18 Neotetracus sinensis
+0.07 Crocidura indochinensis
+0.07 Blarina brevicauda
-0.93 Blarinella quadraticauda
+2.48 Sorex palustris
+0.03 Sorex bedfordiae
-0.46 Sorex araneus
+1.07 Anourosorex squamipes
+3.07 Neomys fodiens
-0.37 Episoriculus fumidus
+0.07 Episoriculus macrurus
+0.07 Episoriculus umbrinus
+2.07 Nectogale elegans
+2.07 Chimarrogale platycephala
+2.07 Chimarrogale styani
+1.85 Solenodon paradoxus
-0.10 Sus scrofa
+1.67 Equus caballus
+0.87 Felis catus
+1.18 Pteropus alecto
+0.18 Cavia porcellus
K98N
D126N
G5D
L9K
L40R
K113T
Q116K
A125D
D53G
Q128K
G129E
N12H
E27A
Q113K
A117E
N35D
L40R
Q116K
Q128E
D35N
Q26R
N35D
D44A
Q113K
S132K
Q116K
S117E
E27L
E27A
R31S
R31S
D53G
H48N
G5E, N12K
E41Q, Q81E
S117D, G124D
D126K, Q128K
G129D
L9H, E27A
K34L, N35D
E54A, Q116K
S117E
(A1V)
(H24Y)
D122N
S132K
E136Q
G35D
Q116K
D53A
G5D
G35D
D53A
Q113K
Q116K
A125D
D126N
K34Q
H48N
E27A
Q113D
Q116K
N81D
(G1A)
D53G
H119Q
Q119H
N12D, A27E
E38Q, E52V
K102E, E109K
G121D, D126S
Q128K, K132E
N140H
S132K
N12K
D53A
E27A
E27A
R31S
A57C
Q116K
A120E
D122A
G124D
D126K
P37K
K132N
G35D
Q116K
A125D
S51K
Q113E
G121D
D125A
D126N
G129D
D44S
E83Q
Q81H
A87K
E83A
K87T
K34T, D53A, Q113H, Q116H
R31S, Q113H, G121H, S132R
S132K
+1.07
+0.07
+0.18
-0.93
-0.46
+0.90
-0.82
+2.07
100
80
60
40
20
0
Time before present (Ma)

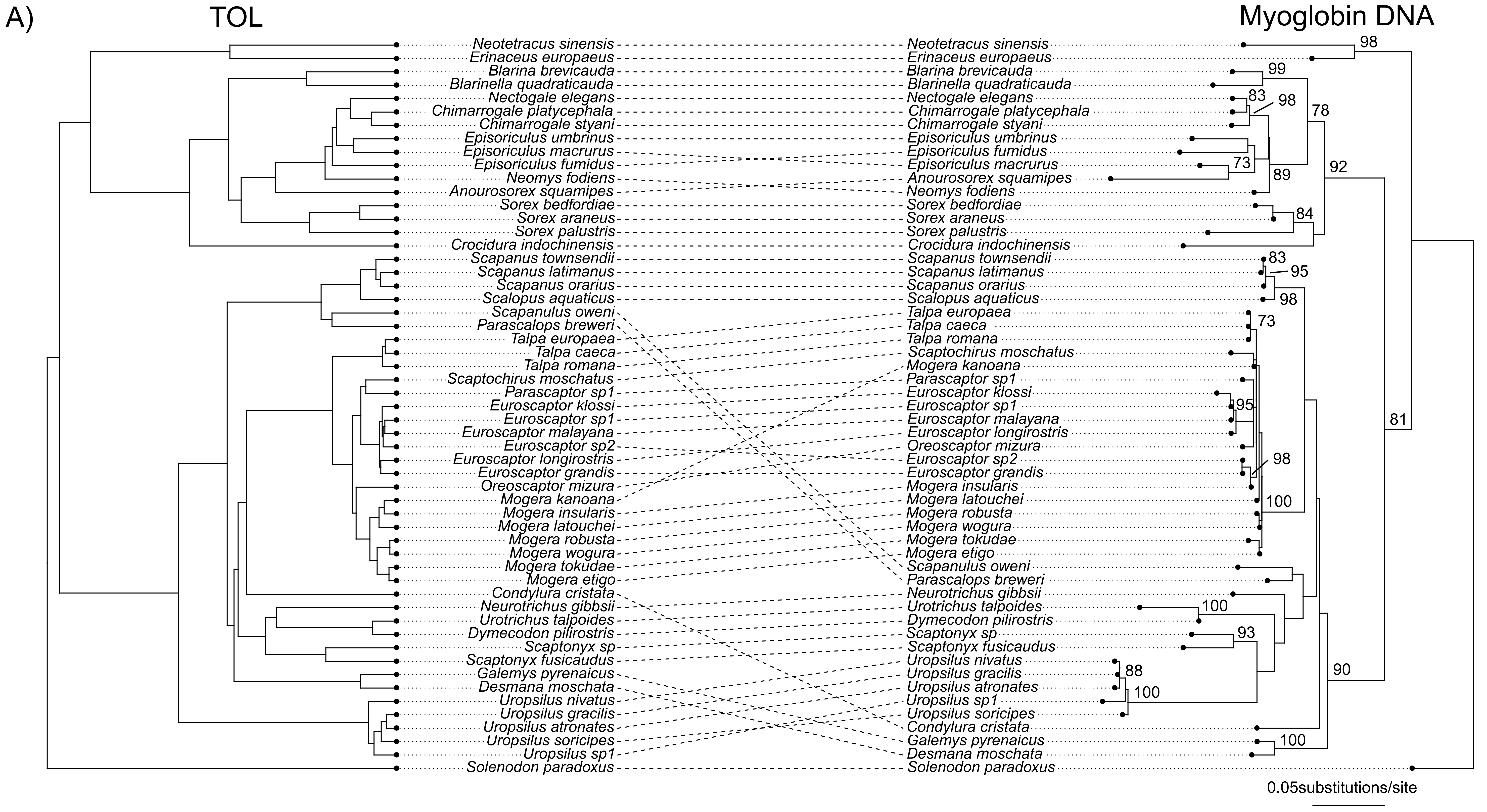
A)
TOL
Myoglobin DNA
Neotetracus sinensis
Erinaceus europaeus
Blarina brevicauda
Blarinella quadraticauda
Nectogale elegans
Chimarrogale platycephala
Chimarrogale styani
Episoriculus umbrinus
Episoriculus macrurus
Episoriculus fumidus
Neomys fodiens
Anourosorex squamipes
Sorex bedfordiae
Sorex araneus
Sorex palustris
Crocidura indochinensis
Scapanus townsendii
Scapanus latimanus
Scapanus orarius
Scalopus aquaticus
Scapanulus oweni
Parascalops breweri
Talpa europaea
Talpa caeca
Talpa romana
Scaptochirus moschatus
Parascaptor sp1
Euroscaptor klossi
Euroscaptor sp1
Euroscaptor malayana
Euroscaptor sp2
Euroscaptor longirostris
Euroscaptor grandis
Oreoscaptor mizura
Mogera kanoana
Mogera insularis
Mogera latouchei
Mogera robusta
Mogera wogura
Mogera tokudae
Mogera etigo
Condylura cristata
Neurotrichus gibbsii
Urotrichus talpoides
Dymecodon pilirostris
Scaptonyx sp
Scaptonyx fusicaudus
Galemys pyrenaicus
Desmana moschata
Uropsilus nivatus
Uropsilus gracilis
Uropsilus atronates
Uropsilus soricipes
Uropsilus sp1
Solenodon paradoxus
98
99
83
98
78
73
89
92
84
83
95
98
73
95
98
100
81
100
93
88
100
90
100
0.05substitutions/site

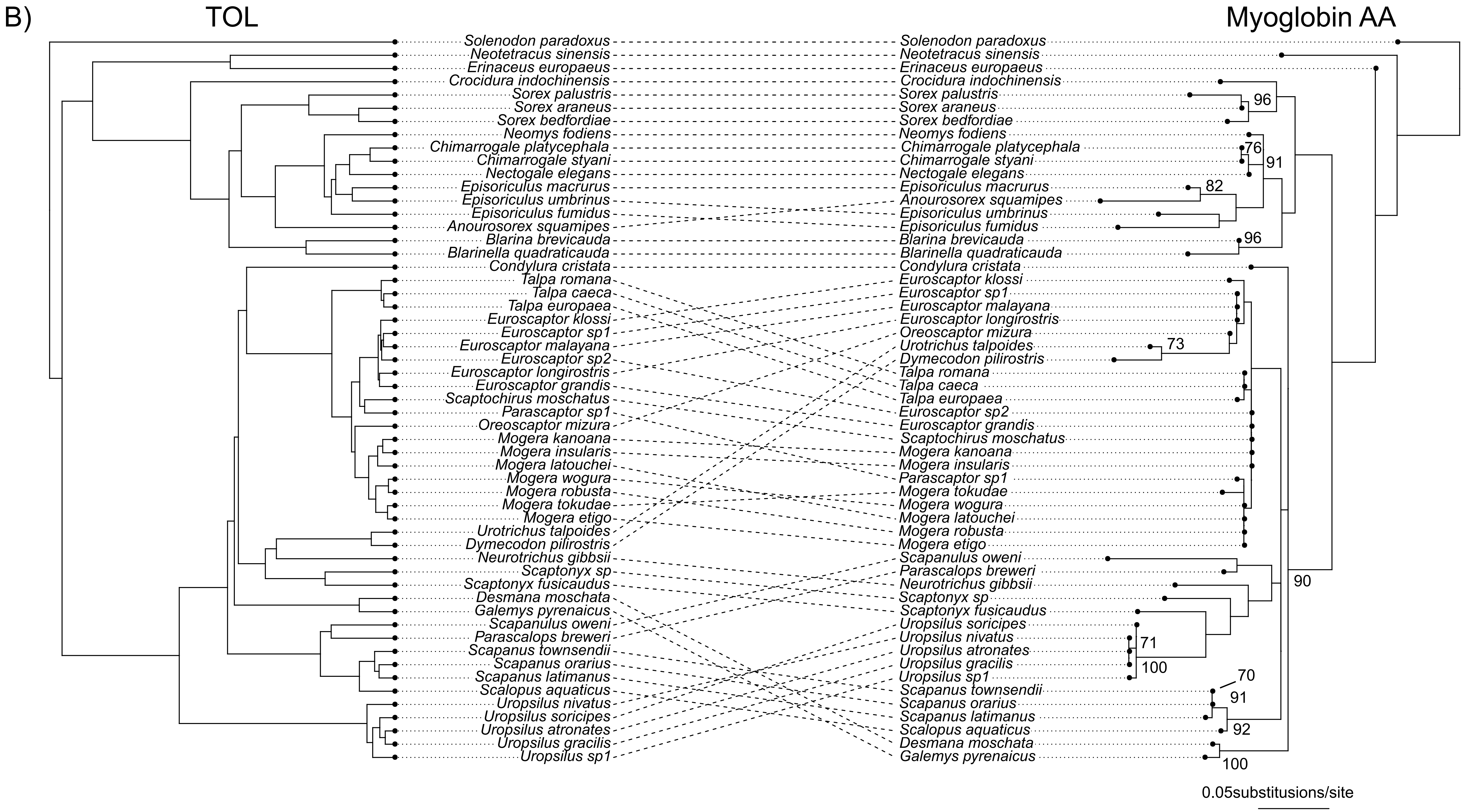
B)
TOL
Myoglobin AA
Solenodon paradoxus
Neotetracus sinensis
Erinaceus europaeus
Crocidura indochinensis
Sorex palustris
Sorex araneus
Sorex bedfordiae
Neomys fodiens
Chimarrogale platycephala
Chimarrogale styani
Nectogale elegans
Episoriculus macrurus
Episoriculus umbrinus
Episoriculus fumidus
Anourosorex squamipes
Blarina brevicauda
Blarinella quadraticauda
Condylura cristata
Talpa romana
Talpa caeca
Talpa europaea
Euroscaptor klossi
Euroscaptor sp1
Euroscaptor malayana
Euroscaptor sp2
Euroscaptor longirostris
Euroscaptor grandis
Scaptochirus moschatus
Parascaptor sp1
Oreoscaptor mizura
Mogera kanoana
Mogera insularis
Mogera latouchei
Mogera wogura
Mogera robusta
Mogera tokudae
Mogera etigo
Urotrichus talpoides
Dymecodon pilirostris
Neurotrichus gibbsii
Scaptonyx sp
Scaptonyx fusicaudus
Desmana moschata
Galemys pyrenaicus
Scapanulus oweni
Parascalops breweri
Scapanus townsendii
Scapanus orarius
Scapanus latimanus
Scalopus aquaticus
Uropsilus nivatus
Uropsilus soricipes
Uropsilus atronates
Uropsilus gracilis
Uropsilus sp1
96
76
91
82
96
73
90
71
100
70
91
92
100
0.05substitusions/site

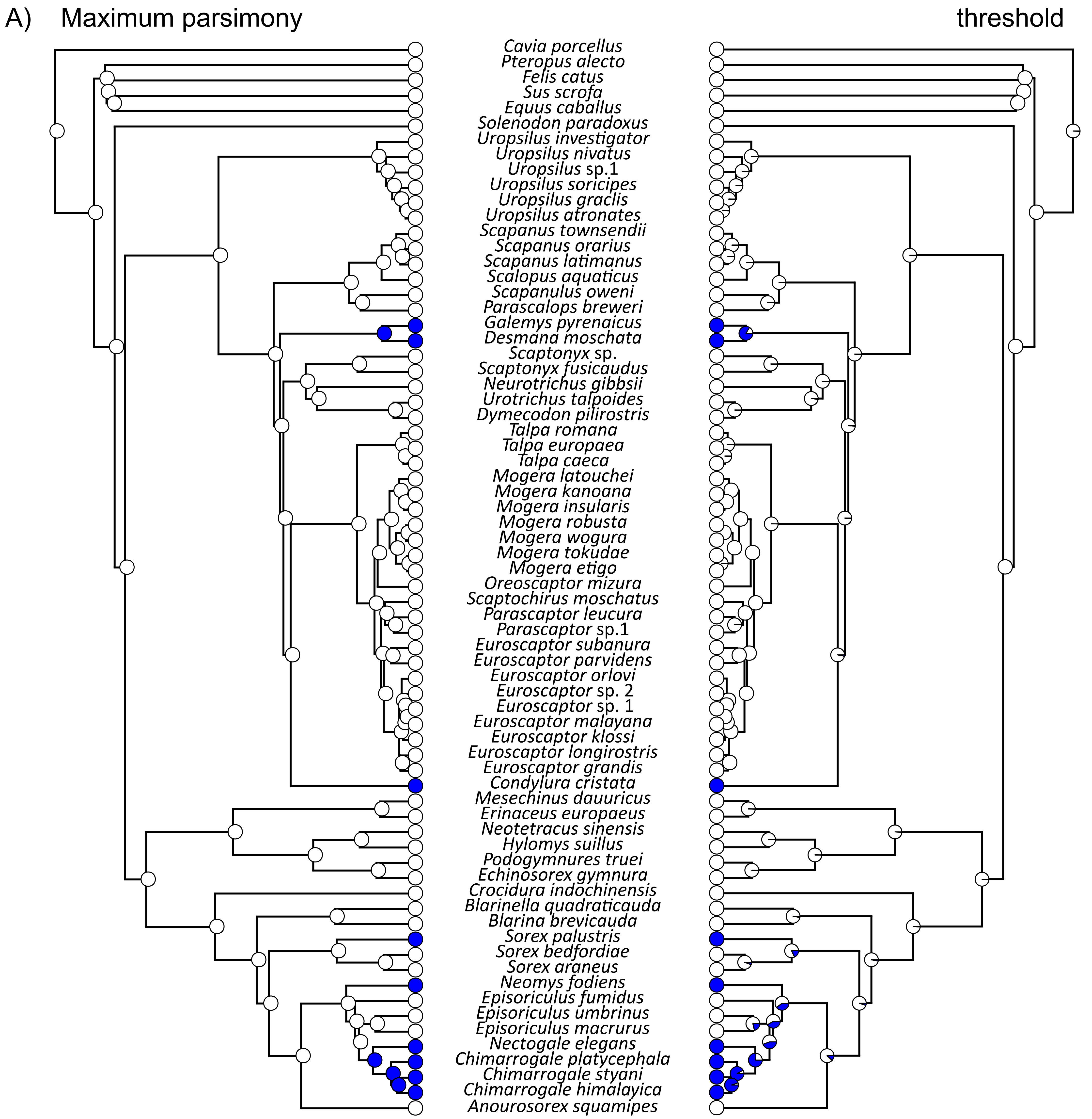
A)
Maximum parsimony
threshold
Cavia porcellus
Pteropus alecto
Felis catus
Sus scrofa
Equus caballus
Solenodon paradoxus
Uropsilus investigator
Uropsilus nivatus
Uropsilus sp.1
Uropsilus soricipes
Uropsilus graclis
Uropsilus atronates
Scapanus townsendii
Scapanus orarius
Scapanus latimanus
Scalopus aquaticus
Scapanulus oweni
Parascalops breweri
Galemys pyrenaicus
Desmana moschata
Scaptonyx sp.
Scaptonyx fusicaudus
Neurotrichus gibbsii
Urotrichus talpoides
Dymecodon pilirostris
Talpa romana
Talpa europaea
Talpa caeca
Mogera latouchei
Mogera kanoana
Mogera insularis
Mogera robusta
Mogera wogura
Mogera tokudae
Mogera etigo
Oreoscaptor mizura
Scaptochirus moschatus
Parascaptor leucura
Parascaptor sp.1
Euroscaptor subanura
Euroscaptor parvidens
Euroscaptor orlovi
Euroscaptor sp. 2
Euroscaptor sp. 1
Euroscaptor malayana
Euroscaptor klossi
Euroscaptor longirostris
Euroscaptor grandis
Condylura cristata
Mesechinus dauuricus
Erinaceus europaeus
Neotetracus sinensis
Hylomys suillus
Podogymnures truei
Echinosorex gymnura
Crocidura indochinensis
Blarinella quadraticauda
Blarina brevicauda
Sorex palustris
Sorex bedfordiae
Sorex araneus
Neomys fodiens
Episoriculus fumidus
Episoriculus umbrinus
Episoriculus macrurus
Nectogale elegans
Chimarrogale platycephala
Chimarrogale styani
Chimarrogale himalayica
Anourosorex squamipes

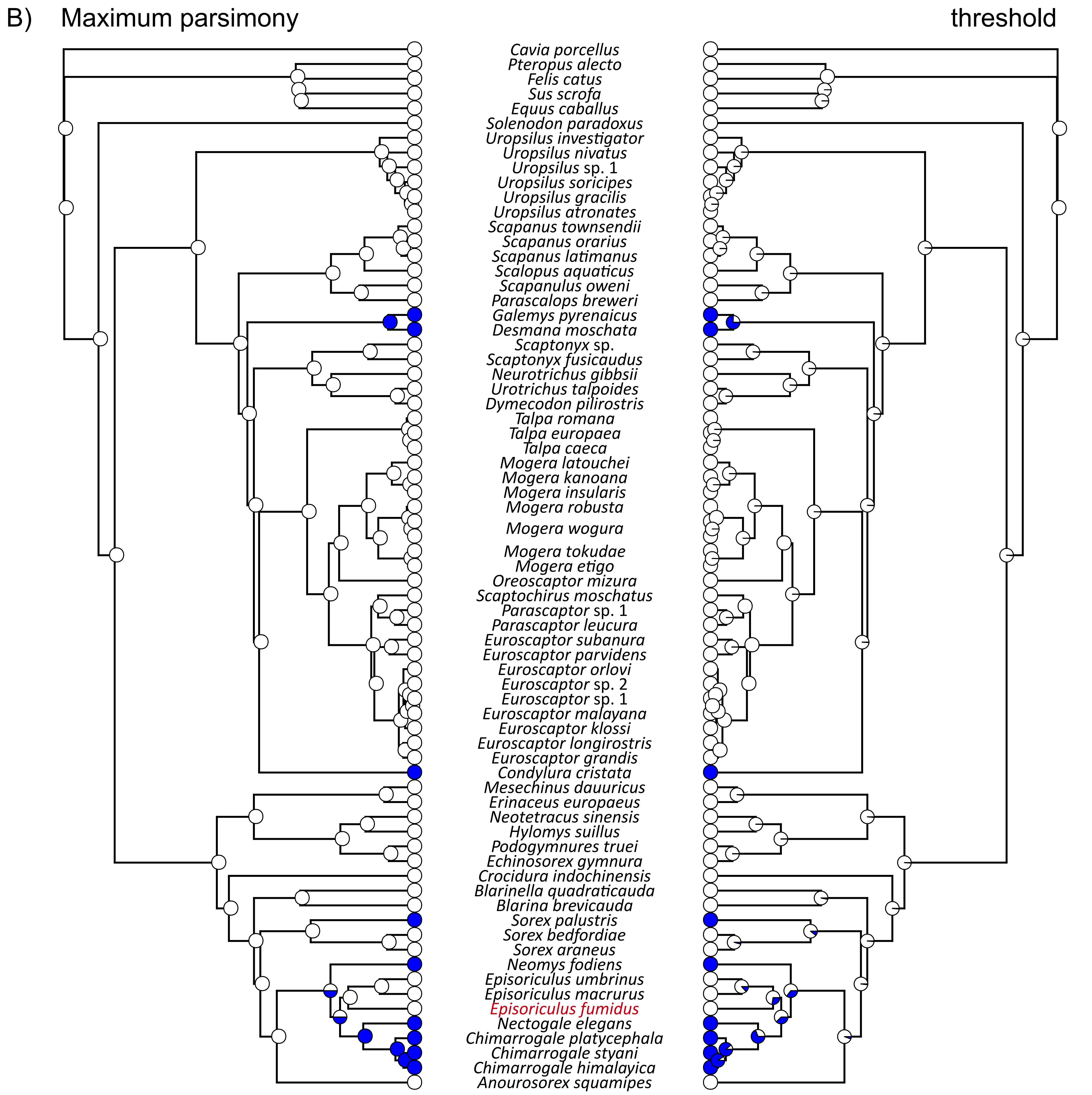

B)
Maximum parsimony
threshold
Cavia porcellus
Pteropus alecto
Felis catus
Sus scrofa
Equus caballus
Solenodon paradoxus
Uropsilus investigator
Uropsilus nivatus
Uropsilus sp. 1
Uropsilus soricipes
Uropsilus gracilis
Uropsilus atronates
Scapanus townsendii
Scapanus orarius
Scapanus latimanus
Scalopus aquaticus
Scapanulus oweni
Parascalops breweri
Galemys pyrenaicus
Desmana moschata
Scaptonyx sp.
Scaptonyx fusicaudus
Neurotrichus gibbsii
Urotrichus talpoides
Dymecodon pilirostris
Talpa romana
Talpa europaea
Talpa caeca
Mogera latouchei
Mogera kanoana
Mogera insularis
Mogera robusta
Mogera wogura
Mogera tokudae
Mogera etigo
Oreoscaptor mizura
Scaptochirus moschatus
Parascaptor sp. 1
Parascaptor leucura
Euroscaptor subanura
Euroscaptor parvidens
Euroscaptor orlovi
Euroscaptor sp. 2
Euroscaptor sp. 1
Euroscaptor malayana
Euroscaptor klossi
Euroscaptor longirostris
Euroscaptor grandis
Condylura cristata
Mesechinus dauuricus
Erinaceus europaeus
Neotetracus sinensis
Hylomys suillus
Podogymnures truei
Echinosorex gymnura
Crocidura indochinensis
Blarinella quadraticauda
Blarina brevicauda
Sorex palustris
Sorex bedfordiae
Sorex araneus
Neomys fodiens
Episoriculus umbrinus
Episoriculus macrurus
Episoriculus fumidus
Nectogale elegans
Chimarrogale platycephala
Chimarrogale styani
Chimarrogale himalayica
Anourosorex squamipes

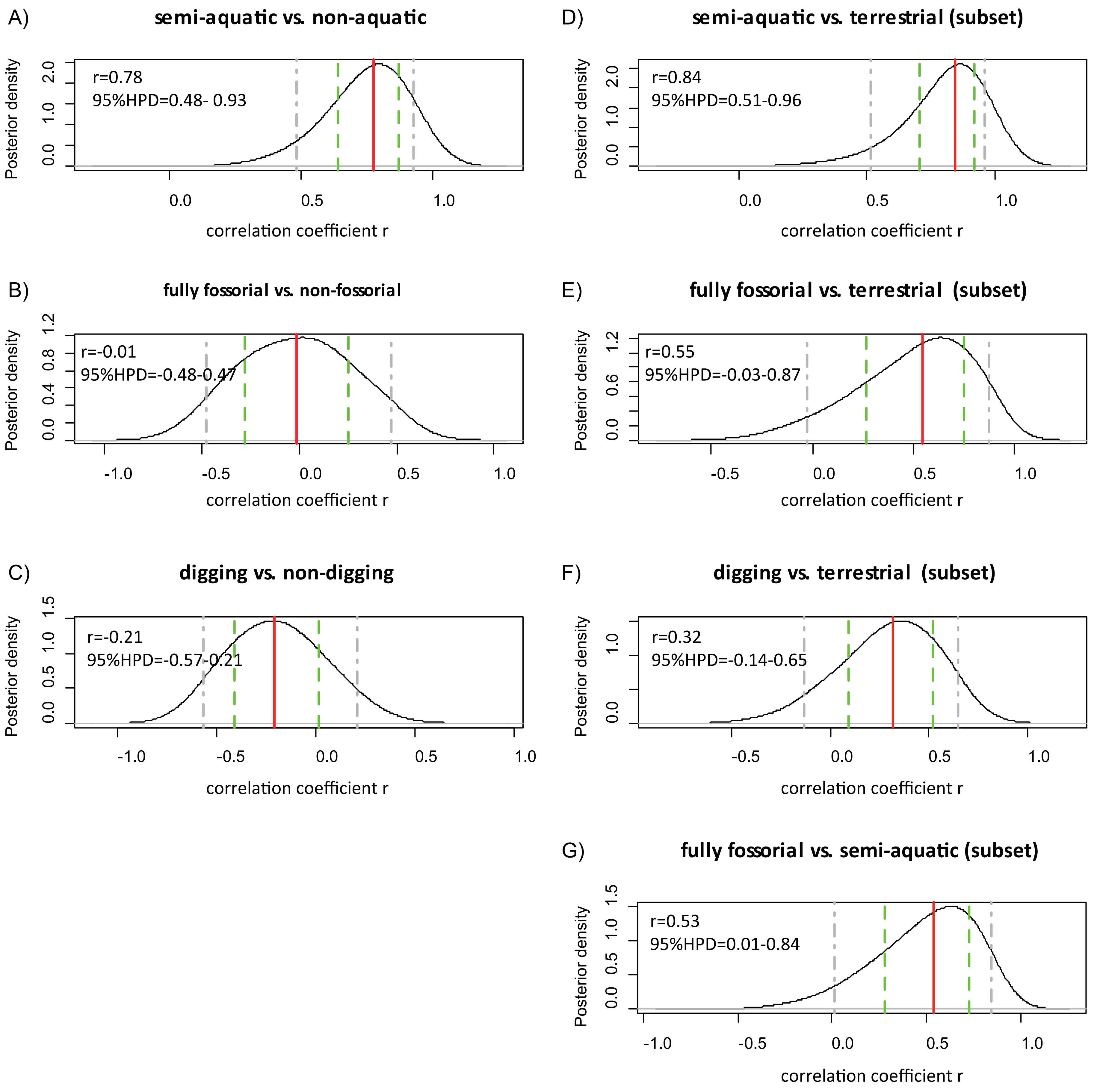
A)
semi-aquatic vs. non-aquatic
r=0.78
95%HPD=0.48- 0.93
Posterior density
correlation coefficient r
B)
fully fossorial vs. non-fossorial
r=-0.01
95%HPD=-0.48-0.47
Posterior density
correlation coefficient r
C)
digging vs. non-digging
r=-0.21
95%HPD=-0.57-0.21
Posterior density
correlation coefficient r
D)
semi-aquatic vs. terrestrial (subset)
r=0.84
95%HPD=0.51-0.96
Posterior density
correlation coefficient r
E)
fully fossorial vs. terrestrial (subset)
r=0.55
95%HPD=-0.03-0.87
Posterior density
correlation coefficient r
F)
digging vs. terrestrial (subset)
r=0.32
95%HPD=-0.14-0.65
Posterior density
correlation coefficient r
G)
fully fossorial vs. semi-aquatic (subset)
r=0.53
95%HPD=0.01-0.84
Posterior density
correlation coefficient r

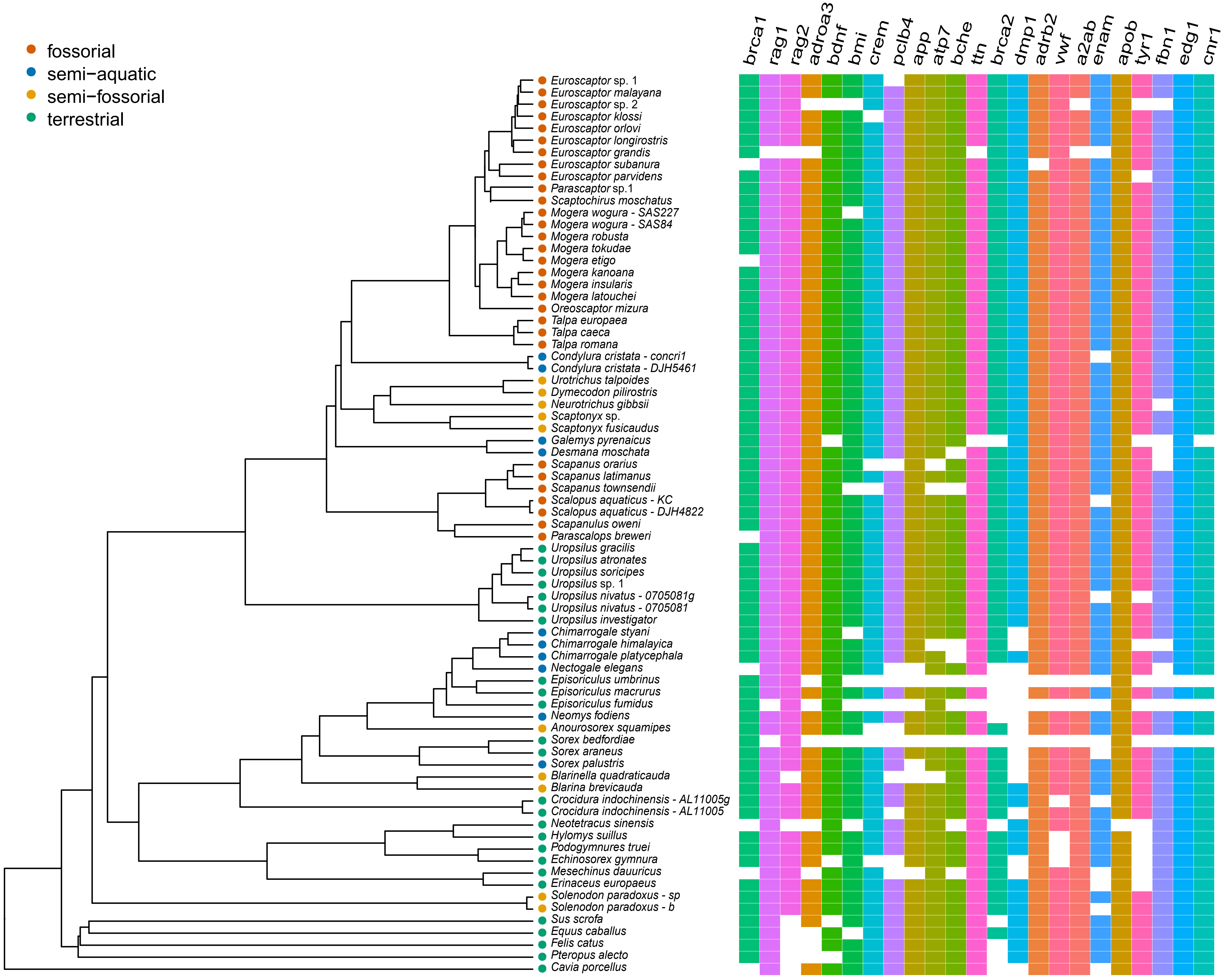

fossorial
semi−aquatic
semi−fossorial
terrestrial
brca1
rag1
rag2
adroa3
bdnf
bmi
crem
pclb4
app
atp7
bche
ttn
brca2
dmp1
adrb2
vwf
a2ab
enam
apob
tyr1
fbn1
edg1
cnr1
Euroscaptor sp. 1
Euroscaptor malayana
Euroscaptor sp. 2
Euroscaptor klossi
Euroscaptor orlovi
Euroscaptor longirostris
Euroscaptor grandis
Euroscaptor subanura
Euroscaptor parvidens
Parascaptor sp.1
Scaptochirus moschatus
Mogera wogura - SAS227
Mogera wogura - SAS84
Mogera robusta
Mogera tokudae
Mogera etigo
Mogera kanoana
Mogera insularis
Mogera latouchei
Oreoscaptor mizura
Talpa europaea
Talpa caeca
Talpa romana
Condylura cristata - concri1
Condylura cristata - DJH5461
Urotrichus talpoides
Dymecodon pilirostris
Neurotrichus gibbsii
Scaptonyx sp.
Scaptonyx fusicaudus
Galemys pyrenaicus
Desmana moschata
Scapanus orarius
Scapanus latimanus
Scapanus townsendii
Scalopus aquaticus - KC
Scalopus aquaticus - DJH4822
Scapanulus oweni
Parascalops breweri
Uropsilus gracilis
Uropsilus atronates
Uropsilus soricipes
Uropsilus sp. 1
Uropsilus nivatus - 0705081g
Uropsilus nivatus - 0705081
Uropsilus investigator
Chimarrogale styani
Chimarrogale himalayica
Chimarrogale platycephala
Nectogale elegans
Episoriculus umbrinus
Episoriculus macrurus
Episoriculus fumidus
Neomys fodiens
Anourosorex squamipes
Sorex bedfordiae
Sorex araneus
Sorex palustris
Blarinella quadraticauda
Blarina brevicauda
Crocidura indochinensis - AL11005g
Crocidura indochinensis - AL11005
Neotetracus sinensis
Hylomys suillus
Podogymnures truei
Echinosorex gymnura
Mesechinus dauuricus
Erinaceus europaeus
Solenodon paradoxus - sp
Solenodon paradoxus - b
Sus scrofa
Equus caballus
Felis catus
Pteropus alecto
Cavia porcellus
